# E/I ratio and net E+I strength are differentially affected across brain disorders

**DOI:** 10.1101/2025.08.15.670484

**Authors:** Arthur-Ervin Avramiea, Marina Diachenko, Andrew Westbrook, Richard Hardstone, Huibert D. Mansvelder, Hilgo Bruining, Klaus Linkenkaer-Hansen

## Abstract

Neuronal oscillations wax and wane near criticality, switching between high- and low-power states. However, the link between this bistability and excitation–inhibition balance is not well understood. Using a computational model, we demonstrate that bistability is influenced by two key factors: the excitatory-inhibitory (E/I) ratio and the combined strength of excitatory and inhibitory activity (E+I). We developed algorithms to infer both metrics from electrophysiological data and validated them in vivo. In intracranial recordings from epilepsy patients, both E/I and E+I increased during seizures, while interictally, E+I remained elevated in seizure zones as E/I decreased—consistent with compensatory inhibition. Extending the framework to scalp EEG from eight disorders, we found that Alzheimer’s disease is better classified by E/I changes, whereas autism spectrum disorder and ADHD are distinguished by E+I alterations. Our findings highlight the importance of bistability in understanding EI regulation and offer handles for diagnostic and treatment decision support.

## Introduction

The delicate balance between excitation (E) and inhibition (I) in neural networks is the result of a complex interplay of regulatory mechanisms that unfold throughout an individual’s development^1–4^. Studies of neurodevelopmental and neurodegenerative disorders indicate that when this equilibrium is not achieved or sustained, deviations in E/I balance can drive circuit-level dysfunction^5–7^. However, the considerable variability in symptom presentation and the underlying neural circuit alterations among individuals—even those with the same disorder^8,9^—emphasizes the need to identify the circuit-level deficits as a basis for targeted interventions.

Research has often concentrated on excitatory/inhibitory (E/I) ratio, but theory suggests that the overall synaptic drive—reflected by the combined strength of E and I (E+I)—is also important. By increasing both E and I, an inhibition-stabilized network can enter a strong-coupling regime, which enhances response gain and alters oscillatory dynamics even if the ratio remains close to one^10,11^. Because no non-invasive marker of net E+I strength exists, its role in human neurological conditions remains an open question.

Recent theoretical advancements infer the E/I ratio from local field potentials^9,12^. These methods leverage the observation that balanced excitation and inhibition tune a network to the critical point—the boundary between ordered and disordered activity—where fluctuations in oscillatory activity from large neuronal populations acquire scale-free statistics^13^. Even modest departures from balanced E/I drive the network away from the critical point, and the resulting loss of scale-free structure serves as an indirect marker of this imbalance. Such shifts are accompanied by diminished functions normally supported by criticality, including the ability to integrate information across distant areas in the brain^14^, as well as to filter and distinguish stimuli^15^. Deviations from the critical-state signatures have been reported across neurodevelopmental disorders such as schizophrenia^16^ and autism spectrum disorder (ASD)^9,17^, as well as across neurodegenerative disorders such as Alzheimer’s^18^ and Parkinson’s^19^, making criticality a useful framework for studying the profile of E/I alterations observed in these disorders.

Intriguingly, networks operating near the critical point are also characterized by spontaneous transitions between a high- and a low-powered state of oscillations^20^, a phenomenon termed spectral bistability, because the system toggles between two stable activity regimes. These transitions are pervasive during waking rest, particularly in the alpha band^20–22^. Preliminary evidence suggests that bistability is influenced by conditions with assumed alterations in E/I ratio^9,13^, such as those induced by GABA-ergic anesthetics ^23^ or epileptogenic networks^22^. Yet, the physiological and computational connections between bistability, E/I ratio and E+I strength are unknown, despite its potential to reveal new aspects of network organization and function.

We hypothesize that bistability provides a framework for assessing the EI network states, both in terms of E/I ratio and net E+I strength. To examine this, we used a computational model where criticality in both neuronal avalanches and neuronal oscillations depend on a balance between excitatory and inhibitory connectivity^13,15^. We first confirmed that, in this model, the presence of scale-free (critical) oscillations co-occurs with bistable switching of oscillations between low- and high-power states^24^. We then found that the prevalence of high-power states in bistable networks can be used to infer E/I ratio, whereas the separation between high- and low-power states can be used to estimate the net strength of excitatory and inhibitory connections (E+I). Finally, we validated these bistability-derived biomarkers in intracranial EEG from epilepsy patients and in scalp EEG recordings from eight neurological or psychiatric disorders, showing differential contribution of E/I and E+I across disorders.

## Results

### Bistability co-occurs with critical oscillations in silico

To study the impact of E/I ratio on bistability, we used the critical oscillations (CROS) model^9,13,15^, which consists of 75% excitatory and 25% inhibitory integrate-and-fire neurons placed on a 50 × 50 grid (**Fig. S1A**). Each neuron connects to a given percentage of neurons in its local range, and by separately modulating the percentage of connected neighbors of excitatory and inhibitory neurons, we can determine the excitatory/inhibitory (E/I) ratio. Varying these parameters reveals that the criticality of neuronal avalanches (**Fig. 1A)** and the criticality of neuronal oscillations (**Fig. S1C**) co-exist for networks with balanced excitatory and inhibitory connectivity. Previously^20^, it was found in a Kuramoto model and in empirical resting-state data, that critical oscillations also exhibit bistability in power of oscillations. Here, we improve on a previously used bistability index (BIS) to make it insensitive to signal length^20,21,25^ (see *Materials and Methods*, **Fig. S2**) and show that bistability indeed co-emerges with criticality of oscillations also in the CROS model (**Fig. S1C–E**). In doing so, we focused our analysis on the alpha (8–16 Hz) band, where CROS shows increasing narrow-band oscillation power as a function of E/I ratio (**Fig. S1B**).

**Figure 1.**
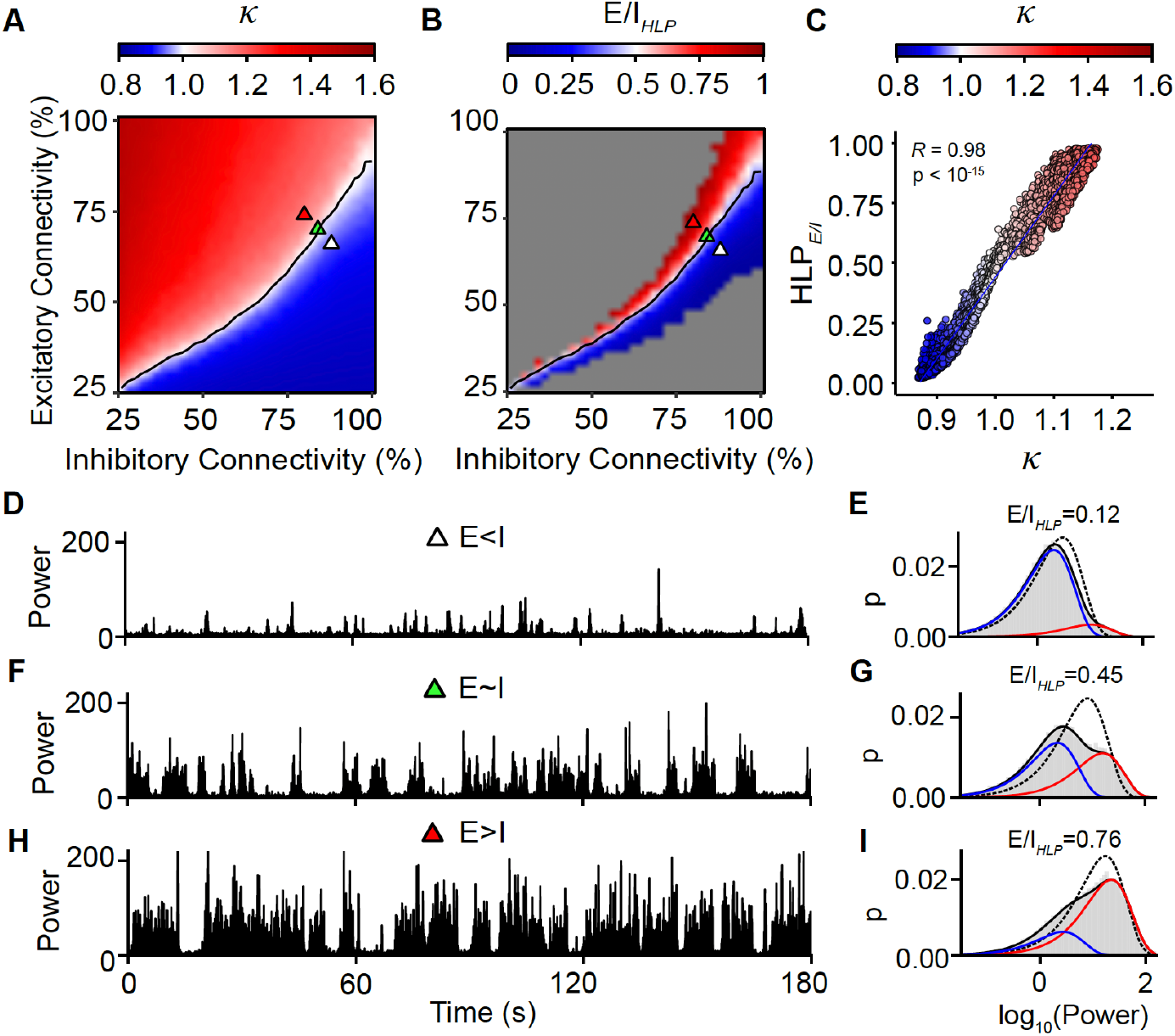
Proportion of High- and Low-power oscillations reflects E/I ratio in critical networks. **(A)** Criticality of avalanches depends on excitation inhibition balance in the CROS model, as reflected by the kappa indices below and above 1 for sub- and super-critical dynamics, respectively. **(B)** Proportion of high- and low-power oscillations (HLP) in the alpha-band (8–16 Hz) follows a similar pattern and **(C)** correlates strongly with Kappa (each dot represents an individual simulation), allowing it to infer E/I ratio. For this reason, we refer to this measure as E/I_*HLP*_. The *gray* area in (**B**) corresponds to networks where E/I_*HLP*_ was not computed, either because the networks did not show enough evidence of bistability (BiS < 2.5) or because E/I_*HLP*_ is very close to 0 (E/I_*HLP*_ < 0.02) or 1 (E/I_*HLP*_ > 0.98). (**D**–**I**) Example traces and distributions illustrate how network criticality affects E/I_*HLP*_. (**D**) Inhibition-dominated networks (*blue* triangle in B) are marked by long periods of relative silence in oscillatory activity, interrupted by brief oscillatory bursts, which is reflected in (**E**) a larger low-power peak (*blue*) and smaller high-power peak (*red*), resulting in an E/I_*HLP*_<0.5 (*black* continuous curve represents the bi-exponential fit of the oscillation-power distribution (*light gray*); the *black* dashed line represents the mono-exponential fit of the data. In contrast, excitation-dominated networks are marked by long oscillatory bursts, interrupted by brief periods of quiescence (**H**, *red* triangle in **B**), which is reflected in a larger high-power peak compared to the low-power peak (E/I_*HLP*_ > 0.5, **I**). In critical networks, there is a balanced amount of quiet and bursty activity (**F**, *green* triangle in **B**), reflected in an E/I_*HLP*_ of ~0.5 (**G**). Overall, the bi-exponential fit comes much closer to the data than the mono-exponential fit.

### The proportion of high- and low-power oscillations in bistable networks predicts E/I in silico

It has been reported that the emergence and duration of UP states in neuronal firing is regulated by the E/I ratio^26,27^. We hypothesized that the emergence of high-power states is also influenced by E/I. To investigate this in the CROS model, we picked three sample networks with increasing E/I ratios, orthogonal to the critical line (**Fig. 1A**) to understand the impact of E/I on bistability. As in Bruining *et al*.^9^, we used the K_*size*_ of avalanches as a proxy for functional E/I ratio. This approach minimizes the influence of random variations in network initialization which are irrelevant to the population activity, and focuses instead on the effective emergent dynamics, with K_*size*_<1 reflecting inhibition-dominated networks, K_*size*_>1 excitation-dominated networks, and K_*size*_~1 balanced networks. The time series of the amplitude envelope of alpha-band oscillations reveals a continuous alternation between low- and high-power states, with the latter being more prevalent in excitation-dominated networks (**Fig. 1D,F,H**). Indeed, bistable network oscillations are characterized by two distinguishable power peaks, where the low-power peak corresponds to the quiescent oscillatory phase and the high-power peak corresponds to the active oscillatory phase (**Fig. 1E,G,I**). Fitting a bi-exponential model to the log_10_ oscillatory power distribution reveals that the exponential associated with the high-power state increases in height, while the exponential associated with the low-power state decreases, as E/I ratio is increased.

We use these E/I dependent shifts in the proportion of high- and low-power oscillations as the basis for developing a new E/I_*HLP*_ biomarker that takes a value close to 0 for networks devoid of high-power oscillatory activity, and 1 for networks with sustained high-power oscillations (*Materials and Methods*). Since the fitting of exponentials corresponding to high- and low-power states is unreliable for networks with little evidence of bistability, based on the computation model, networks with BIS<2.5, as well as networks with E/I_*HLP*_<0.02 or E/I_*HLP*_>0.98 were excluded from the computation of E/I_*HLP*_ (**Fig. S3A**,**B**). After applying this selection criterion, we found that as the structural E/I is gradually increased, networks spend less time in the low-powered state and more time in the high-powered state, reflected in ever-increasing values of E/I_*HLP*_ (**Fig. 1B,E,G,I**). Close to the critical region, E/I_*HLP*_ is close to 0.5, reflecting a balanced contribution of low- and high-powered state to the overall network dynamics for networks with balanced E/I. The shifts in E/I_*HLP*_ following changes in E/I ratio can also be observed across the spectrum in narrow frequency bins (**Fig. S4**), consistent with previous reports in Diachenko et *al*.^12^ for other biomarkers relating E/I to criticality. Importantly, compared to the previously developed fE/I method^9^, E/I_*HLP*_ shows a superior ability to distinguish between excitation-dominated, balanced, and inhibition-dominated networks (**Fig. 1C**, compared with **Fig. S5B**), while also tracking synaptic gain changes akin to those induced by neuromodulation^28^ (**Fig. S6A–C**).

### Increasing net E+I strength separates oscillation power peaks

Typically, alterations in excitation and inhibition are studied through the lens of their ratio. However, it is possible to imagine scenarios in which both excitatory and inhibitory activity increase, but the E/I ratio stays the same. To the best of our knowledge, there is at present no algorithm that can non-invasively infer net E+I strength from population activity. One way to conceptualize net E+I strength structurally, is as the total number of excitatory and inhibitory connections realized in a network, from the total possible, which we coin connection density (**Fig. 2A**).

**Figure 2.**
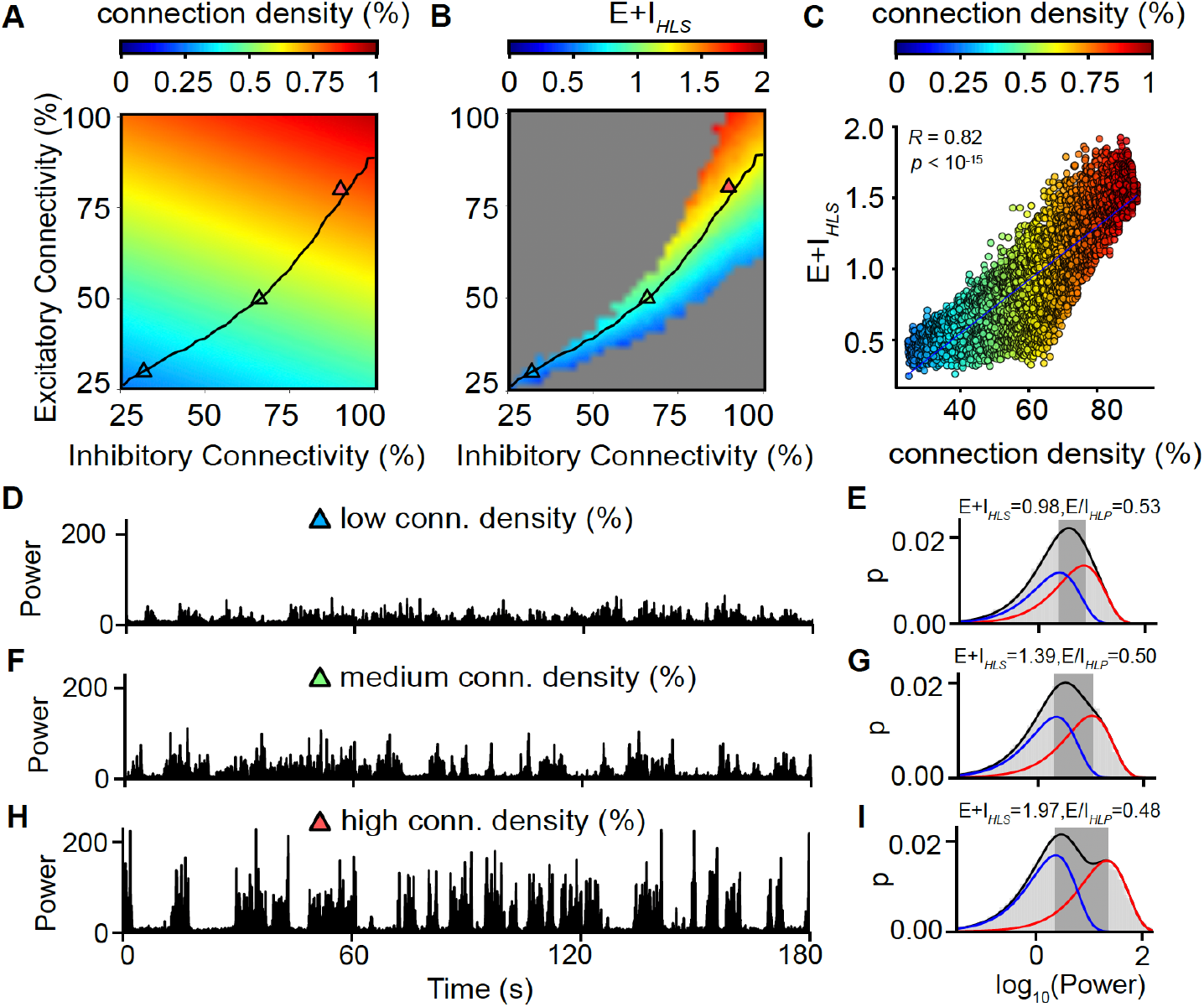
Increasing density of functional connections (E+I) causes a separation of oscillation power peaks. (**A)** We computed connectivity density as the proportion of synapses present in a network relative to the total number of possible synapses (critical networks are indicated by the *black* line). Connectivity density increases when either excitatory connectivity % or inhibitory connectivity % goes up. (**B**,**C**) The separation between the high- and low-power peaks (E+I_*HLS*_) increases with increasing connectivity density (each dot in **C** represents an individual simulation). For this reason, we refer to this measure as E+I_*HLS*_. The *gray* area in (**B**) corresponds to networks where E/I_*HLP*_ was not computed, either because the networks did not show enough evidence of bistability (BiS < 2.5) or because E/I_*HLP*_ is very close to 0 (E/I_*HLP*_ < 0.02) or 1 (E/I_*HLP*_ > 0.98) (**D**-**I**) Example traces and bimodal distributions illustrate how network connectivity density affects the separation of high- and low-power states. We picked three networks from the critical line (**A**) and observed that as we increase the connectivity density, oscillation power bursts stand out more relative to quiet intervals. This is reflected in an increased distance between the high- and low-power peaks (**E**,**G**,**I**) *dark*-*gray* rectangle; the *black* continuous curve represents the bi-exponential fit of the oscillation power distribution, comprised by a low-power peak (*blue* curve) and a high-power peak (*red* curve). The histogram of the data is represented with *light gray*.

In the CROS model, bistability is explained by two features: the proportion of high- and low-power states, and the separation between high- and low-power peaks (**Fig. 2B,E,G,I**). The proportion of high-to-low power accounts for ~50% of variance in bistability (quadratic model, *adjusted R*^*2*^*=0*.*52*) whereas the separation of high- and low-power peaks also accounts for ~50% (quadratic model, *adjusted R*^*2*^*=0*.*53*). Together they account for ~95% of the variance in bistability (*adjusted R*^*2*^*=0*.*95*). Since we have already established that E/I_*HLP*_ is a strong predictor of E/I, we next evaluated, across the phase space of the CROS model, whether the separation of high- and low-power peaks (E+I_*HLS*_) is dependent on the connection density (**Fig. 2B**). Since the estimation of high- and low-power peaks is unreliable for networks with low evidence of bistability, we applied a similar threshold as for E/I_*HLP*_, excluding networks with BIS<2.5, E/I_*HLP*_<0.02 or E/I_*HLP*_>0.98. We found it to be strongly correlated with connection density (**Fig. 2C**). Namely, for networks with the same level of E/I (close to the critical line), those that have a higher connection density exhibit more pronounced transitions between the low- and high-power states, which translate into increased separation between the high- and low-power peaks (**Fig. 2D-I**). The association between connection density and E+I_*HLS*_ (**Fig. 2C**) was markedly stronger than that of connection density with any of the other biomarkers (**Fig. S7**). The shifts in E+I_*HLS*_ following changes in E+I ratio can also be observed across the spectrum in narrow frequency bins (**Fig. S8**). Importantly, the E+I_*HLS*_ is sensitive not only to changes in structural E+I, but also to co-alterations in the synaptic gain of excitatory and inhibitory connections akin to those induced by neuromodulation ^28^ (**Fig. S9**). Thus, we created a novel biomarker that gives us a functional estimate of the net E+I strength, resulting from the long-term structure or from transient changes (e.g., neuromodulation).

### Ictal periods exhibit elevated E/I ratio and net E+I strength compared to inter-ictal periods

Epileptiform activities are widely recognized as extreme cases of E/I imbalance. Perturbations of synaptic dynamics that increase the E/I ratio can trigger seizures^29,30^, whereas perturbations that decrease the E/I ratio can stop ongoing seizures^31,32^. We analyzed pre-surgical intracranial EEG recordings from 58 subjects with drug resistant epilepsy^33^. Biomarker values were computed and averaged separately for ictal and interictal periods for each subject and across good channels belonging to the seizure onset zone (SOZ) and channels outside of this region (nSOZ) (see *Materials and Methods*). In both SOZ and nSOZ channels, E/I_*HLP*_ as well as E+I_*HLS*_ were significantly higher during the ictal compared to the interictal periods across the entire frequency spectrum (paired *t*-test, FDR corrected with *q* = 0.05) (**Fig. 3A–D**). We zoomed in on one frequency band (10.5–13.4 Hz) and visualized the results for one representative subject, for a channel closest to the center of the SOZ region (**Fig. 3E–H**), and a nSOZ channel furthest away from SOZ (**Fig. 3I–L**). The shifts in E/I_*HLP*_ and E+I_*HLS*_ could easily be traced back to the time series of oscillatory power, with the prevalence of high-power oscillations, and the separation between high- and low-power states clearly increasing in the ictal state. This indicates that both E/I_*HLP*_ and E+I_*HLS*_ can track profound E/I alterations in occurring between ictal and interictal periods.

**Figure 3.**
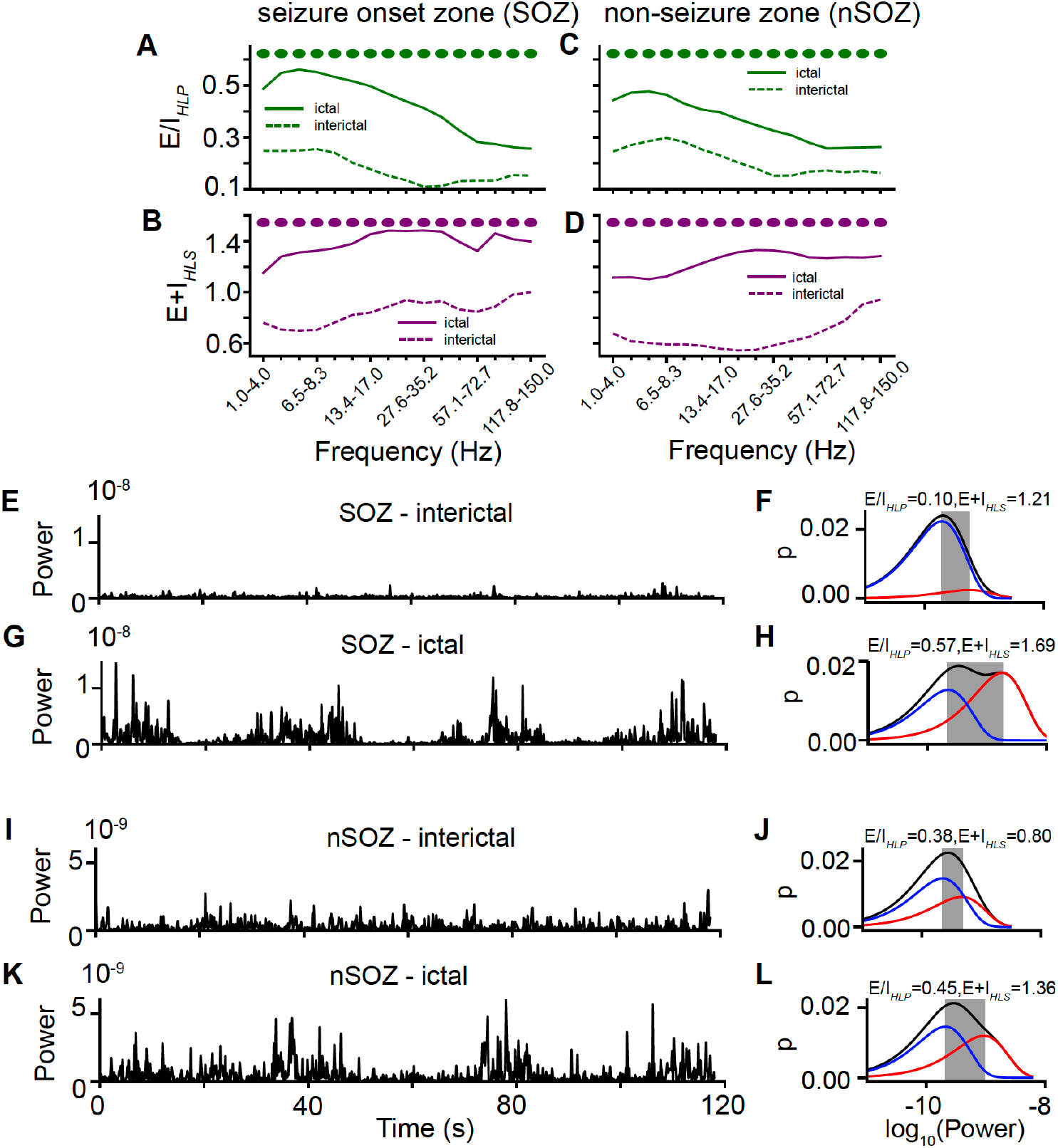
E/I ratio and net E+I strength are elevated during ictal periods. Both seizure zone (**A**,**B**,**E–H**) and non-seizure zone (**C**,**D**,**I-L**) electrodes show increases in E/I_*HLP*_ and E+I_*HLS*_ during ictal periods. Biomarker values were averaged across seizure (SOZ)/non-seizure zone (nSOZ) electrodes, and a paired *t*-test was run across subject-level averages. Circles (**A–D**) correspond to frequency bins where comparisons stayed significant after FDR correction (*q*=0.05). (**E–L**) A representative subject was selected, and sample traces of the oscillatory power time series were plotted for a selected frequency band (10.5-13.4 Hz), along with the bimodal power distributions, to illustrate the changes in oscillatory dynamics between the ictal and interictal states.

### E/I ratio reflects inhibitory suppression of seizure zone during interictal periods

During interictal periods, it has been hypothesized that SOZ is actively suppressed by the surrounding regions^34–36^. Indeed, we found that the SOZ showed significantly reduced E/I_*HLP*_ in the entire 5.1–150 Hz range when compared to nSOZ during the interictal period (paired *t*-test, FDR corrected with *q* = 0.05) (**Fig. 4A**), in agreement with the inhibitory suppression hypothesis. Nonetheless, E+I_*HLS*_ remained significantly higher in SOZ compared to nSOZ during the interictal period (**Fig. 4B**). During the ictal period, the SOZ showed significantly higher E/I_*HLP*_ in the 1–73 Hz range, and significantly higher E+I_*HLS*_ in the 4–17 Hz range, when compared to the nSOZ (**Fig. 4C,D**). We zoomed in on one frequency band (10.5–13.4 Hz) and visualized the results for one representative subject, for a channel closest to the center of the SOZ region (**Fig. 4E–H**), and a nSOZ channel furthest away from SOZ (**Fig. 4I-L**). Although the effects were more subtle when comparing SOZ vs. nSOZ activity within interictal/ictal periods are more subtle than the changes between interictal and ictal periods, they could still be detected by E/I_*HLP*_ and E+I_*HLS*_, suggesting they could be clinically important biomarkers

**Figure 4.**
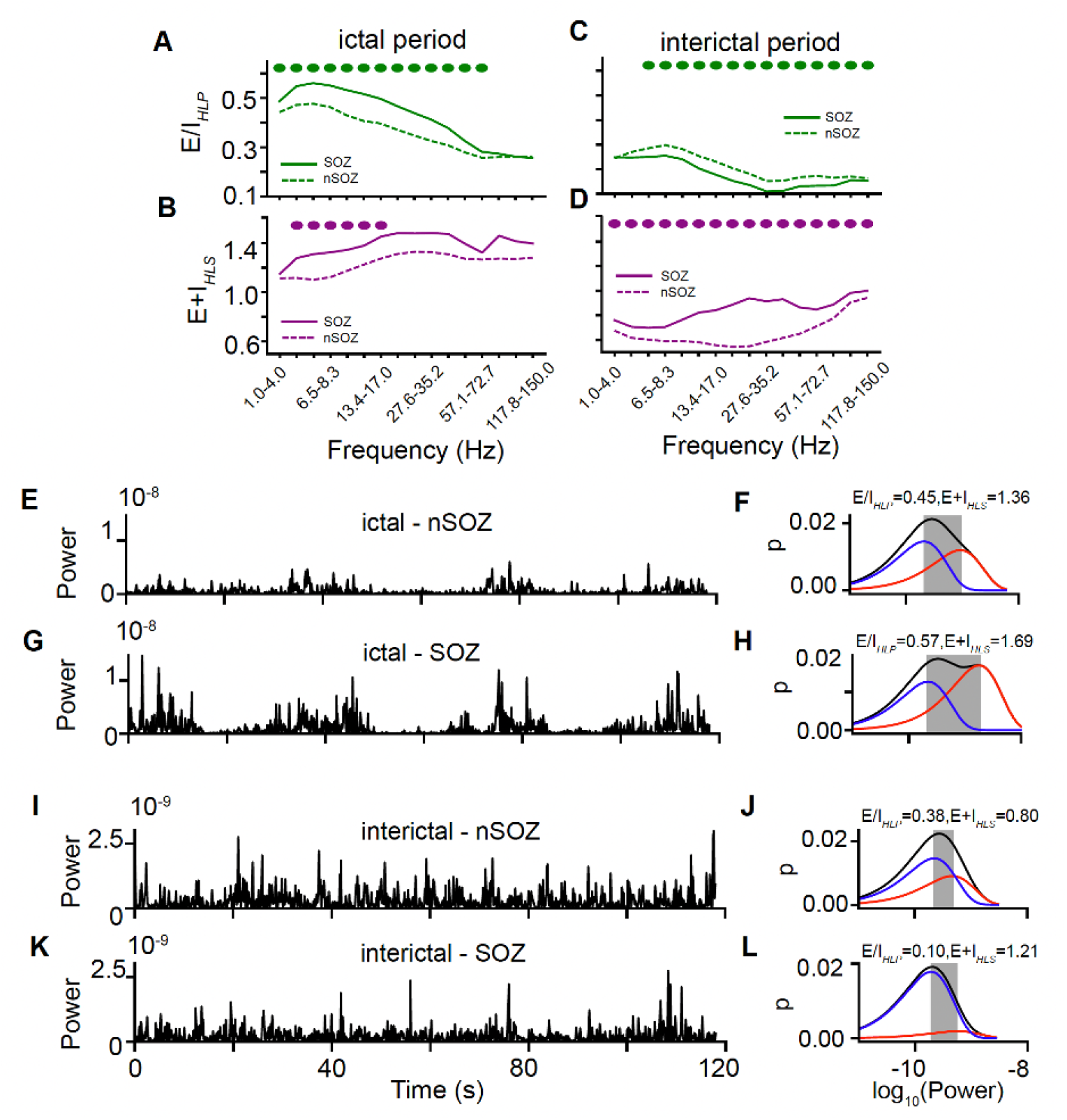
E/I ratio reflects seizure zone suppression during interictal intervals. **(A)** During ictal periods, the seizure zone (SOZ) exhibits significantly higher E/I_*HLP*_ compared to non-seizure zone (nSOZ). **(C)** During interictal periods, on the other hand, the seizure zone has lower E/I_*HLP*_. **(B**,**D)** E+I_*HLS*_ is significantly higher in the seizure zone regardless of the seizure status. Biomarker values were averaged across seizure (SOZ)/non-seizure zone (nSOZ) electrodes, and a paired *t*-test was run across subject-level averages. Circles (**A–D**) correspond to frequency bins where comparisons stayed significant after FDR correction (*q* = 0.05). (**E–L**) A representative subject was selected, and sample traces of the oscillatory power time series were plotted for a selected frequency band (10.5-13.4 Hz), along with the bimodal power distributions, to illustrate differences in oscillatory dynamics between SOZ and nSOZ.

### Brain disorders show disrupted E/I ratio and net E+I strength

Having determined that E/I_*HLP*_ and E+I_*HLS*_ can track epilepsy-related changes in excitatory and inhibitory dynamics, our next goal was to test the ability of these biomarkers to differentiate healthy controls from individuals with various brain conditions. For this purpose, we used resting-state EEG from cohorts with Alzheimer’s disease, attention deficit and hyperactivity disorder (ADHD), autism spectrum disorder (ASD), insomnia, major depressive disorder (MDD), obsessive compulsive disorder (OCD), Parkinson’s disorder, and tinnitus. We employed machine learning to determine the individual and combined contributions of E/I_*HLP*_ and E+I_*HLS*_ in distinguishing a disordered brain state from a healthy one. We computed the biomarker values and then averaged across electrodes to get a whole-brain average for each subject and for each frequency bin (see *Materials and Methods*), resulting in a data vector per subject with length of 22 (11 frequency bins for 2 biomarkers: E/I_*HLP*_ and E+I_*HLS*_). Each disordered group was paired with a healthy control group. We employed logistic regression with elastic net regularization for feature selection, which combines both L1 (lasso) and L2 (ridge) penalties to balance feature sparsity and multicollinearity. Model evaluation was conducted using repeated nested cross-validation with five outer folds (see *Materials and Methods*). In brief, for each fold, an elastic-net logistic regression model was trained on the remaining four folds, with optimal hyperparameters selected via grid-search cross-validation. Model’s performance was evaluated by calculating the Area Under the Receiver Operating Characteristic Curve (AUC-ROC) for each fold. To mitigate split bias, the entire procedure was repeated ten times. AUC-ROC scores from the resulting 50 outer folds were then averaged to provide a robust estimate of the model’s predictive accuracy. Performance was benchmarked against a null AUC-ROC distribution generated by repeating the same procedure on data with randomly shuffled class labels using 500 permutations.

The disorders lay on a spectrum where Alzheimer’s disease, ADHD, and ASD are well distinguished from healthy controls by alterations in E/I_*HLP*_ and E+I_*HLS*_, whereas other disorders (e.g., MDD and tinnitus) showed weak, if any, discrimination based on changes in excitatory inhibitory dynamics (**Fig. 5A**).

**Figure 5.**
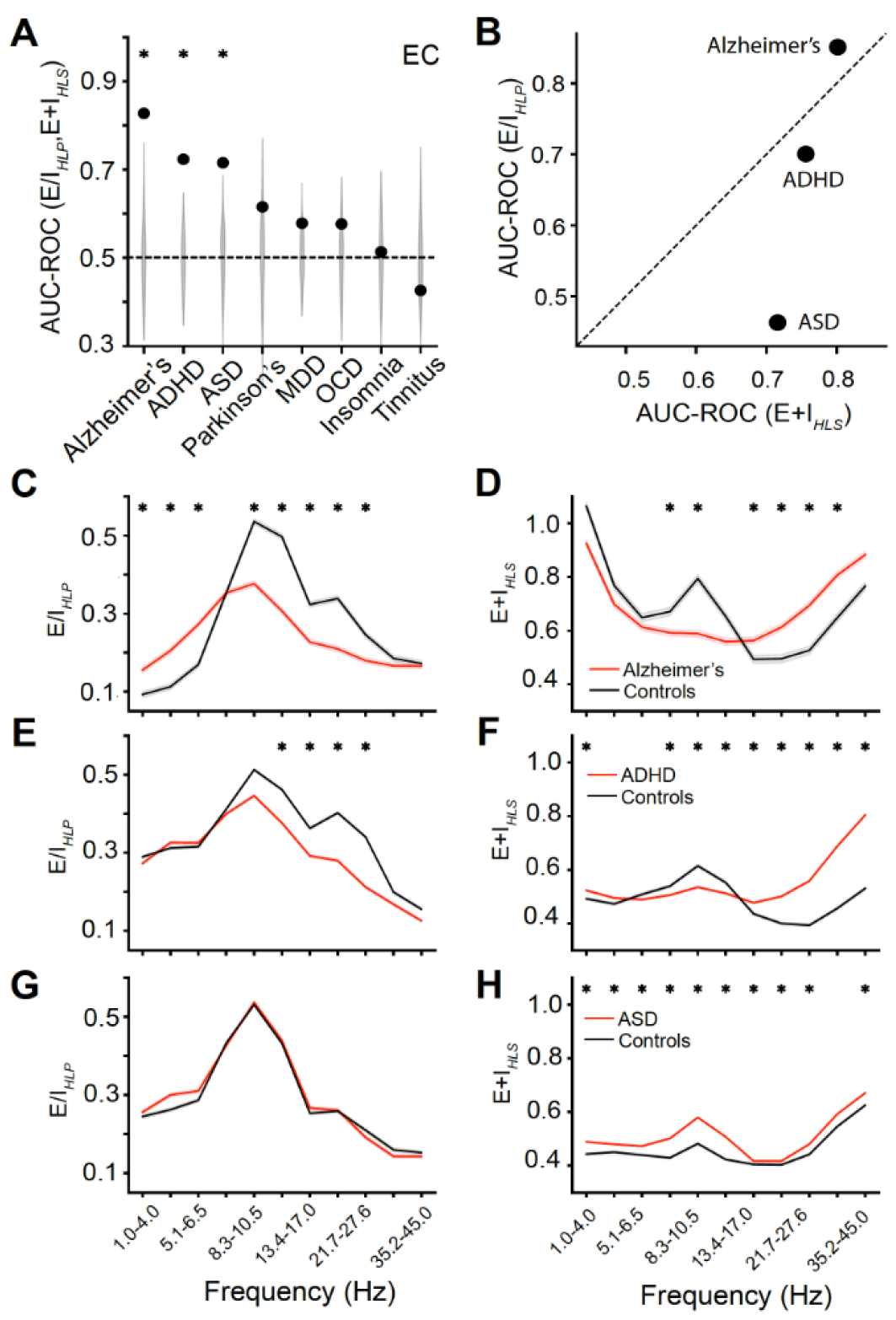
Disrupted E/I ratio and E+I density across brain disorders. **(A)** Logistic regression models reveal that patients with Alzheimer’s, ADHD, or ASD can be distinguished from healthy controls by alterations in E/I_*HLP*_ and E+I_*HLS*_. Significance for each disorder was assessed by comparing the average AUC-ROC (black dot) across 50 cross-validation folds against a null distribution generated with 500 random permutations (gray violin plot). Black asterisk reflects significance after FDR (*q* = 0.05) correction. **(B)** When training logistic regression separately for E/I_*HLP*_ and E+I_*HLS*_, we found Alzheimer’s disease showing mostly disruptions in E/I ratio, while ASD and ADHD more prominent changes in E+I density. To illustrate how the relative contribution of E/I_*HLP*_ and E+I_*HLS*_ differs across the three disorders, we plotted disorder-specific case-control averages of each biomarker across the frequency spectrum (**C–H**). Asterisks above a trace mark frequency bins in which patient and control means differ significantly (two-sample *t*-test, FDR-corrected at *q* = 0.05, performed separately for every disorder-by-biomarker combination).

When evaluated separately, E/I_*HLP*_ demonstrated higher classification accuracy than fE/I^6^—a previously established method for E/I estimation—in Alzheimer’s and ADHD (**Fig. S10A**). For the remaining disorders, neither model achieved significant classification performance. Additionally, E/I_*HLP*_ was applicable to a larger percentage of signals (**Fig. S10B**).

To determine whether Alzheimer’s disease, ADHD, and ASD show preferential alterations in either E/I_*HLP*_ or E+I_*HLS*_, we re-trained using only the E/I_*HLP*_ data, or only the E+I_*HLS*_ data. For all disorders, paired comparisons of fold accuracy (AUC-ROC, 50 folds) between the two models were significant, even after correction for multiple comparisons (*t*-test, FDR correction with *q* = 0.05, **Table S1**). Alzheimer’s disease was better distinguished from healthy controls by alterations in E/I ratio (**Fig. 5B–D**), while ASD and ADHD subjects were better discriminated by alterations in net E+I strength (**Fig. 5B,E–H**). This suggests that the dual dimensions of excitatory-inhibitory dynamics are useful to understand alterations occurring in disorders, with some disorders better understood through the lens of one or the other dimension of EI.

## Discussion

Using a computational model, we determined that two features of the bistability of neuronal oscillations can index complementary dimensions of excitatory-inhibitory interactions: the proportion of high- and low-power oscillations reflects E/I ratio (E/I_*HLP*_), whereas the separation of high- and low-power oscillations tracks net E+I strength (E+I_*HLS*_). In epilepsy—a paradigmatic condition of pathologically elevated excitation—both biomarkers proved sensitive: during ictal periods E/I_*HLP*_ and E+I_*HLS*_ rose across all measured areas. Between seizures, however, the seizure focus showed a divergent pattern: E+I_*HLS*_ remained elevated while E/I_*HLP*_ dropped below values in the adjacent non-seizure zone, consistent with the interictal suppression hypothesis. Subsequently, by leveraging machine learning, we evaluated the utility of the two bistability-based biomarkers, E/I ratio and net E+I strength, in differentiating brain disorders from the healthy state. We found that while E/I ratio alterations predominantly drove classification accuracy in Alzheimer’s disease, net E+I strength alterations contributed more to differentiation of ASD and ADHD from controls. This distinction highlights specific contributions of E/I versus E+I disorder-specific patterns in neuronal dynamics to brain disorders, with the potential to guide future mechanism-focused investigations of pathophysiology and treatment.

Our findings extend previous work linking higher E/I ratio to increased prevalence of UP states by showing a similar effect on the proportion of high- and low-power oscillatory states, revealing a shared basis in excitatory-inhibitory interactions. These studies have shown that various mechanisms, including adaptation^37,38^, synaptic depression^39^, and balance of excitatory and inhibitory synapses^40^, influence UP state proportion. While the specifics of these mechanisms vary, they can be interpreted as functional alterations in the E/I ratio, with distinct biophysical origins. Indeed, the E/I_*HLP*_ metric can track changes in the E/I ratio attributable to either structural connectivity or synaptic gain.

To assess the performance of our new bistability-based metric, E/I_*HLP*_, we compared it to the established fE/I algorithm^9^, which was successfully applied to track E/I ratio alterations in autism spectrum disorder (ASD)^9,41^, Alzheimer’s^42^, epilepsy^22^ and STXBP1^43^, among other disorders. E/I_*HLP*_ showed clear advantages in both modeling and empirical data. In the computational model, E/I_*HLP*_ tracked E/I ratio with near-perfect correlation to the criticality index κ (*r* = 0.98), outperforming fE/I (*r* = 0.86). In empirical datasets, E/I_*HLP*_ better distinguished patients from controls, particularly in conditions with above-chance classification. In addition, E/I_*HLP*_ offers the advantage of applicability across a greater percentage of signals, making it a robust alternative to fE/I in capturing EI-related brain state alterations. While fE/I and E/I_*HLP*_ are based on different principles—temporal fluctuation structure vs. bistable power ratios—their convergence supports the validity of E/I estimation. Notably, E/I_*HLP*_ improves interpretability by linking directly to observable features of oscillatory dynamics.

Alongside the widely used E/I ratio, the net E+I strength offers an equally important, independent dimension—one that captures shifts in overall synaptic drive and helps distinguish whether observed E/I changes arise from altered excitation, inhibition, or both. This distinction matters because many physiological and pathological processes are invisible to the ratio alone. In disorders where EI balance is disrupted, compensatory mechanisms often engage, sometimes with secondary adverse effects on brain function^44,45^. Using our metrics, we were able to indirectly track EI dynamics in epilepsy. While an increased E/I ratio is often expected in epileptogenic networks, we observed evidence of inhibitory suppression within the seizure onset zone (SOZ), suggesting a compensatory response^34–36^. Despite this suppression, the functional E+I density remains elevated in the SOZ compared to the surrounding areas, reflecting both inhibitory control and latent excitation poised to trigger further seizures. Similar patterns of inhibitory suppression have also been documented in other developmental conditions associated with epileptogenic networks, such as ASD^9,41^, STXBP1^43^, and tuberous sclerosis complex^46^. Applying our bistability-derived metrics to source-reconstructed EEG in future studies could deepen understanding of these compensatory interactions across regions and help clarify its functional consequences.

Using machine learning, we assessed how E/I ratio and net E+I strength differentiated individuals with brain disorders from healthy controls. While prior studies have often attempted to link ASD and ADHD to E/I imbalances^9,47,48^, our findings suggest that net E+I strength provides a more sensitive dimension for distinguishing these groups at the population level. In contrast, Alzheimer’s disease was better differentiated by alterations in the E/I ratio. For the remaining conditions, classification performance did not exceed chance levels, possibly due to smaller effects, greater heterogeneity, or mechanisms beyond the scope of these metrics. These results indicate that the relative relevance of E/I versus E+I metrics varies across disorders, motivating future work linking these dimensions more directly to circuit-level mechanisms.

We based the classification analyses on whole-brain averages to avoid overfitting due to limited sample sizes, potentially obscuring regional dissociations in EI alterations. Global functional properties, such as connectivity density of long-range connections, are influenced by local EI^14,49^ and can better characterized by methods that consider interregional dynamics rather than whole-brain averages. Future work should leverage the E/I_*HLP*_ and E+I_*HLS*_ biomarkers to dissect how local connectivity patterns, reflected by high- and low-power oscillations, relate to global interactions and functional outcomes in both healthy and disordered brains.

Together, our findings establish a bistability-based framework for capturing excitatory–inhibitory dynamics in the human brain. The ability to extract interpretable, reliable biomarkers of both E/I ratio (E/I_*HLP*_) and E+I strength (E+I_*HLS*_) opens a new window into the latent circuit architecture of brain disorders. By revealing distinct EI alterations across conditions and identifying patterns of compensatory suppression, these metrics offer a foundation for future diagnostics, patient stratification, and individualized interventions that target the core of neuronal network dysfunction.

## Materials and Methods

### CRitical OScillations (CROS) model

The CROS model is an *in silico* model that was created by Poil *et al*. ^13^ with the aim of having a spiking neuronal network that produced oscillatory activity with long-range temporal correlations in the oscillation amplitude. In this paper, we use an extended version of this model^9^ with optimized parameters to investigate the dependence of neuronal network activation patterns on structural connectivity and synaptic parameters. CROS models a network of 75% excitatory and 25% inhibitory integrate-and-fire neurons arranged in a50 × 50 open grid. Networks differ in their two connectivity parameters, excitatory and inhibitory connectivity, which are the percentage of other neurons within a local range (a square with width = 7 neurons centered on the presynaptic neuron) that each excitatory and inhibitory neuron connects to, respectively. Connectivity parameters were set between 25–100% at 2% intervals, and 20 different networks were created for each combination of excitatory and inhibitory connectivity – *C*_*E*_ and *C*_*I*_, and run for 1000 seconds. For a full description of the model see Bruining et *al*.^9^. To emulate the effects of neuromodulation, we separately modulated the synaptic gain of excitatory-excitatory and excitatory-inhibitory connections, with either gain ranging between 0.1 and 2, with a step of 0.05. The gain was multiplied with the synaptic weight of the corresponding connections, as in Pfeffer et *al*.^28^. For each combination of excitatory-excitatory and excitatory-inhibitory gain, 5 different networks were created, and simulations were run for 1000 seconds. To estimate the net E+I of the network, we simply added the synaptic gain of the excitatory-excitatory and excitatory-inhibitory connections.

### Neuronal avalanches

To get a measure of the activity dynamics of the neuronal network, we applied neuronal avalanche analysis^9,13,15^. A neuronal avalanche is defined as a period where neurons are spiking above a certain threshold—in our case set to half the median of activity. The size of the avalanche is the number of spikes during this period. We then computed the κ index^13,50^, which calculates the difference between the distribution of our data and a power-law, by calculating the average difference of the cumulative distribution of a power-law function, *P*, (with exponent −1.5 for size and −2.0 for duration) and that of our experimental data, *A*, at 10 equally spaced points on a logarithmic axis (*β*) and adding 1.

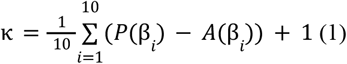

A subcritical distribution is characterized by *κ* < 1, and a supercritical distribution by *κ* > 1, whereas *κ* = 1 indicates a critical network.

### Network activity analysis

A network signal was created by summing the total number of neurons spiking at each time-step with a Gaussian white noise signal of the same length with mean = 0 and σ = 3^9,15^. This level of white noise was set to allow all networks to achieve a time-varying phase, which is not the case without adding the noise, when there are silent periods in the network. Detrended fluctuation analysis, oscillation power, and phase-locking factor, which are described below, were calculated on this pre-processed signal, with Gaussian noise added. The analysis of neuronal avalanches was performed on the raw signal, consisting of the total number of neurons spiking at each time step.

### Epilepsy data

58 subjects with drug resistant epilepsy, at the Hospital of the University of Pennsylvania, underwent intracranial EEG with subdural grid, strip, and depth electrodes (ECoG) or purely stereotactically-placed depth electrodes (SEEG). The dataset is openly available, together with detailed methodology, at Bernabei et *al*.^33^. In brief, each patient also underwent subsequent treatment with surgical resection or laser ablation. Electrophysiologic data for both interictal and ictal periods is available, as are electrode localizations in ICBM152 MNI space. Furthermore, clinically-determined seizure onset channels are provided, as are channels which overlap with the resection/ablation zone, which was rigorously determined by segmenting the resection cavity. Bad channels were marked and excluded from the analysis. Where multiple recordings were available, we concatenated ictal recordings together and interictal recordings together. For the analyses, we included all good channels in the seizure onset area (SOZ) for the SOZ group, and all good channels that did not pertain to either the SOZ or the resection/ablation zone, to the nSOZ group.

### Alzheimer’s disorder data

36 subjects with Alzheimer’s disease and 29 age matched controls, had their EEG measured during resting state eyes-closed. The data is openly available, together with detailed methodology, at Miltiadous et *al*.^51^. In brief, these recordings took place in a clinical routine setting. Recordings were acquired from the 2nd Department of Neurology of AHEPA General Hospital of Thessaloniki by an experienced team of neurologists. A clinical EEG device (Nihon Kohden 2100), with 19 scalp electrodes (Fp1, Fp2, F7, F3, Fz, F4, F8, T3, C3, Cz, C4, T4, T5, P3, Pz, P4, T6, O1, and O2) and 2 electrodes (A1 and A2) placed on the mastoids for an impedance check and as reference electrodes, was used for the recording of the EEG signals. The electrodes were placed according to the 10–20 international system. Each recording was performed according to the clinical protocol with participants being in a sitting position with their eyes closed. The recording montage was referential using Cz for common mode rejection. The sampling rate was 500 Hz and the resolution was 10 uV/mm. The following is the EEG signals’ preprocessing pipeline. The signals were re-referenced to the average value of A1-A2 after applying a Butterworth band-pass filter with a frequency range of 0.5 to 45 Hz. The signals were then subjected to the ASR routine, an automatic artifact reject technique that can eliminate persistent or large-amplitude artifacts, which removed bad data periods that exceeded the maximum acceptable 0.5 s window standard deviation of 17 (which is regarded as a conservative window). The ICA method (RunICA algorithm) was then used to convert the 19 EEG signals to 19 ICA components^52^. ICA components categorized as “eye artifacts” or “jaw artifacts” by the EEGLAB platform’s automatic classification method “ICLabel” were automatically excluded. It should be mentioned that, even though the recording was done in a resting state with the eyes closed, eye movement artifacts were still identified in certain EEG recordings.

### Autism spectrum disorder (ASD) data

100 children with ASD and 29 typically developing children had their EEG measured during resting state eyes-closed. The data has been previously analyzed in ^9,41^, where a full methodology is available. In brief, EEGs were recorded during 3–5 minutes of eyes-closed rest at the UMC Utrecht using A 64-channel BioSemi EEG system at a sampling rate of 2048 Hz and common mode sense (CMS) reference electrode. EEG analyses were initially pre-processed using the in-house developed Neurophysiological Biomarker Toolbox (NBT) written in MATLAB. All recordings were manually cleaned for artifacts, i.e., noisy channels were discarded and noisy intervals removed. Subsequently, the data were re-referenced to the average reference. An average of 217 seconds (range 61–308 s) per recording were available for analysis for the children with ASD and TDC samples.

### Parkinson’s disorder data

We used EEG recordings of 27 Parkinson’s disorder patients (ON medication) and 27 controls, from a study at the University of New Mexico (UNM; Albuquerque, New Mexico). Additionally, we used EEG recordings from OFF medication sessions for the 27 PD patients from UNM which were recorded in the practically defined OFF levodopa period, 12 h after the last dose of dopaminergic medication. Control participants were demographically matched for age and sex with PD patients and did not differ in any measurements of education or premorbid intelligence. Resting state EEG recordings of the UNM subjects were gathered under both eyes-open and eyes-closed conditions. For the OFF medication sessions, the data was gathered only under the eyes-open condition. The data is openly available, together with detailed methodology, at Anjum et *al*.^53^. In brief, EEG was recorded from sintered Ag/AgCl electrodes across 0.1–100 Hz with a sampling rate of 500 Hz on a 64-channel Brain Vision system, with online reference set to channel CPz as baseline. Data were cleaned using an automatic pipeline. First, we downsampled to 250 Hz, and a 1-45 Hz bandpass filter was applied. Then we computed ICA (25 components) using the infomax algorithm, and used ICLabel to automatically mark components. We then removed components marked as ‘eye’ or ‘muscle’ with a >80% probability. Last, we re-referenced the signal of each electrode to the whole-brain average.

### TD-BRAIN data

Eyes-open and eyes-closed EEG were recorded for 200 patients with ADHD, 30 with insomnia, 40 with obsessive compulsive disorder, 319 with major depressive disorder, 30 with Tinnitus, and 47 healthy controls. The dataset is openly available, together with detailed methodology, at van Dijk et *al*.^54^. In brief, 2-minute, 26 channel EEG-recordings were performed, based on the 10–10 electrode international system, using a Compumedics Quickcap or ANT-Neuro Waveguard Cap with sintered Ag/AgCl electrode, acquired at a sampling rate of 500 Hz (low-pass filtered at 100 Hz prior to digitization). Data were assessed during resting state, consisting of: a 2-minute Eyes Open (EO) task, where the subject was asked to rest quietly, with eyes open and focus on the red dot at the center of the computer screen in front of them, and a 2-minute Eyes Closed (EC) task, where the subject was asked to close their eyes and retain the same position as before. The data was preprocessed using a custom python script, available on the project page (www.brainclinics.com/resources).

### Signal pre-processing

First, the signal is filtered in 11 narrow frequency bands for the EEG, spanning 1 to 45 Hz, and 16 narrow frequency bands for the sEEG data, spanning 1 to 150 Hz, with the range of frequency bins optimized in Diachenko et *al*.^12^. We used a finite response filter (FIR), with the default parameters from the MNE (1.7) *filter* function^55^. For the CROS model, we filtered the data in one frequency band: 8 to 16 Hz. For each frequency band, amplitude (A) is extracted using the *apply_hilbert* MNE function. To speed up computation, we downsampled the resulting amplitude to the nearest multiple of 10 above 5 times the upper frequency bound, ensuring a minimum of 5 samples per cycle. Then, power P is calculated as A^2^.

### DFA

The main steps of the DFA algorithm are as follows^12^:

1. The signal is bandpass filtered in the desired frequency range using a finite-impulse-response (FIR) filter.
2. The amplitude envelope of the filtered signal A is extracted using Hilbert transform.
3. Logarithmically spaced window sizes are generated between 0.1 and 1000 seconds, with 20 window sizes for each order of magnitude. The sizes are then converted from seconds to samples using the signal’s sampling frequency.
4. The range of times scales of interest is defined and is called the DFA fitting interval. Frequently, the upper bound is set at 30 seconds, as this is the range within which LRTC are typically reported for human brain oscillations^56^. However, it is also constrained by the length of the signal. To have suitable statistics for the largest window size, one may set it to the signal length divided by 10. When choosing the lower bound of the fitting interval, the integration effect of the underlying filters is considered^12^.
5. For each window size within the fitting range, the signal profile is calculated and split into 50%-overlapping same-sized windows.
6. The mean standard deviation is calculated to obtain the mean fluctuation per window size.
7. The DFA exponent is estimated as the best-fit line of the mean fluctuation as a function of window sizes in log-log coordinates in the DFA fitting interval.

### fE/I

The main steps of the fE/I algorithm are as follows^9,12^:

1. The signal is bandpass filtered in the desired frequency range using a finite-impulse-response (FIR) filter.
2. The amplitude envelope of the filtered signal is extracted using Hilbert transform.
3. The signal profile is calculated and segmented into 80%-overlapping 5-second windows.
4. The windows are normalized using the mean of the amplitude envelope calculated per window.
5. Subsequently, the normalized windows are detrended.
6. The normalized fluctuation function for each window is computed as the root-mean square fluctuation of the detrended amplitude-normalized signal profile.

Finally, the fE/I value is obtained as 1-*r*(*W*_*amp*_,*W*_*nF(t)*_), where *r* is Pearson correlation coefficient and *W*_*amp*_ and *W*_*nF(t)*_ are windowed amplitudes and windowed normalized fluctuations, respectively. The fE/I is set to NaN (i.e., missing) if the computed DFA exponent does not exceed the DFA threshold of 0.6.

### Estimation of the bistability index (BiS)

The BiS index of a power time series (P) is computed from a model comparison between a bimodal or unimodal fit of its probability distribution function (pdf); a large BiS means that the observed pdf is better described as bimodal, and when BiS → 0 the pdf is better described as unimodal. We adapted the approach used in ^25^ to compute BiS. One first modification to the original algorithm is that we normalized the empirical power series P by dividing it to its median, which allows for the BiS index to be effectively estimated for varying orders of magnitude of data. Next, to find the pdf of the normalized power time series, nP was partitioned into 200 equal-distance bins and the number of observations in each bin was tallied. Next, maximum likelihood estimate (MLE) was used to fit a single-exponential function (i.e., the square of a Gaussian process follows an exponential pdf):

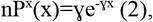

and a bi-exponential function:

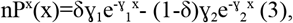

where γ_1_ and γ_2_ are exponents and δ is the weighting factor. The parameter γ in equation (2) is determined through direct computation, and then it is used to initialize optimization of the parameters in equation (3), where the initial value is γ_1=_γ_2_=γ and δ=0.5. Optimization is run using Nelder-Mead optimization, and the parameters γ_1_ and γ_2_ are explored in log space. We constrain the optimization algorithm such that γ_1_>γ_2_, thus γ_1_ refers to the right-most exponential, and γ2 to the left-most exponential.

Next, the Bayesian information criterion (BIC) was computed for the single- and bi-exponential fitting:

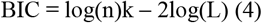

where n is the number of samples; L is the likelihood function; k is the number of parameters: k = 1 for single-exponential BIC_Exp_ and k = 3 for bi-exponential model BIC_biE_. Thus BIC imposes a penalty to model complexity of the bi-exponential model because it has two more degrees of freedom (second exponents and the weight δ) than the single exponential model.

To avoid the likelihood and BIC values from being influenced by the signal length, we made one additional modification to the algorithm in Freyer et *al*.^25^, correcting for the signal length by treating all signals as if they had the length of 100000 samples:

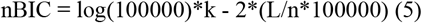

Last, the BiS estimate is computed as the log_10_ transform of difference between the two BIC estimates as

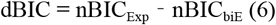

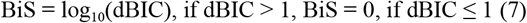

Thus, a better model yields a small BIC value, and BiS will be large if the bi-exponential model is a more likely model for the observed power time series.

### Proportion of High- and Low-Power Oscillations (E/I_*HLP*_)

The constraint for the ordering of γ_1_ and γ_2_ (γ_1_>γ_2_) lends δ to be interpreted as the proportion factor of the two exponentials (equation 3). When δ=0, the left-most exponential is the only one present, and the right-most is absent. When δ=1, the right-most exponential is the only one present, and the left-most is absent. When δ=0.5, both exponentials have equal contributions. Thus, as δ varies from 0 to 1, the contribution of the first peak is reduced, and the contribution of the second peak is increased, with 0.5 corresponding to equal height of the first and second peak.

If we consider, when the bi-exponential model is fit to the histogram of the power data, that the first exponential corresponds to the low-power oscillatory state, and the second exponential corresponds to the high-power oscillatory state, then a δ=0 would correspond to a complete absence of an high-power states, a δ=1 to a complete absence of low-power states, and a δ=0.5 to an equal presence of high- and low-power states.

The bi-exponential parameters are not reliable when fit on data that is not bi-stable. Based on the CROS model, we determine a threshold of BiS>=2.5, over which the δ-BiS relationship is inverse-U shaped (**Fig. S3**), and delta matches tightly with Kappa estimates (**Fig. 1C**). δ values with BiS<2.5 are set to NA. We also set δ to NA, when δ<0.02 or δ>0.98, that is either one of the two peaks in the bi-modal, was very small. We call the δ parameter thus thresholded, the Proportion of high- to low-power oscillations (E/I_*HLP*_):

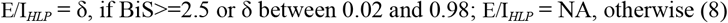

A similar selection criterion operates for fE/I, which cannot be computed for networks with DFA<=0.6, explaining the gray area in **Fig. S5A**.

### Separation of High- and Low-Power Oscillations (E+I_*HLS*_)

Let L be the power value corresponding to the peak of the low-power exponential, and H be the power value corresponding to the peak of the high-power exponential. Then the separation of high- and low-power oscillation can be computed as:

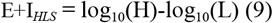

which corresponds to the orders of magnitude separating the high-from the low-power peak. Similar to E/I_*HLP*_, E+I_*HLS*_ values were considered invalid for networks with low evidence of bistability, or with one of the exponentials contributing very little to the total: BIS<2.5, E/I_*HLP*_<0.02 or E/I_*HLP*_>0.98. In these cases, E+I_*HLS*_ was set to NA.

### Machine Learning for Disorder Classification and Biomarker Validation

To assess the ability of the E/I_*HLP*_ and E+I_*HLS*_ biomarkers to distinguish various brain disorders from the healthy state, we used machine learning to compute the Area Under the Receiver Operating Characteristic Curve (AUC-ROC) for each comparison. Each subject’s data consisted of whole-brain averages of E/I_*HLP*_ and E+I_*HLS*_ values across 11 frequency bins, resulting in 22 input features. Where data were missing, it was imputed using the subject’s mean, separately for the E/I_*HLP*_ and E+I_*HLS*_ features. After this, the data were standardized by removing the mean and scaling to unit variance. When evaluating both biomarkers together, all 22 features were used; for single-biomarker analyses, we used only the 11 features related to that biomarker.

We employed logistic regression with elastic net regularization which combines both L1 (lasso) and L2 (ridge) penalties to balance feature sparsity and multicollinearity. To ensure robustness, we implemented a repeated nested cross-validation procedure with five outer and five inner folds, preserving class proportions across all splits. Each outer fold served as an independent test set, while the remaining four folds were used to train the model. Prior to training, optimal hyperparameters were selected via five-fold grid-search cross-validation, repeated five times with different fold splits. The grid spanned the inverse regularization strength C ∈ {0.001, 0.01, 0.1, 1} and the elastic-net mixing parameter l1-ratio ∈ {0, 0.25, 0.5}. To mitigate split bias, the entire nested cross-validation procedure was repeated ten times, resulting in 50 outer test folds. Model performance was evaluated using AUC-ROC for each outer fold. Final performance was reported as the mean AUC-ROC across all test folds. Statistical significance was assessed by comparing the observed mean AUC-ROC to a null distribution generated from 500 permutations of randomly shuffled class labels, repeating the same nested cross-validation procedure. Comparison of accuracy between the E/I_*HLP*_ and E+I_*HLS*_ models for disorders with significant model performance on the combined E/I_*HLP*_ and E+I_*HLS*_ features was performed using paired *t*-test on the fold-based AUC-ROC scores computed for each model across 50 folds. FDR correction with *q* = 0.05 was used to correct for multiple comparisons across disorders.

When comparing E/I_*HLP*_ with fE/I, we used the same training procedure, taking into account only the 11 frequency bin values for either of the biomarkers. Comparison of accuracy between the E/I_*HLP*_ and fE/I models was performed using paired *t*-test on the fold-based AUC-ROC scores computed for each model across 50 folds, but only for disorders where at least one model showed significantly above-chance AUC-ROC compared to its respective permutation-based null distribution. FDR correction with *q* = 0.05 was used to correct for multiple comparisons across disorders.

## Funding

Amsterdam UMC TKI grant BRAINinBALANCE, project number 31556-2012377 (HB, KLH) MGH FMD ECOR Clinical Research Fellowship (2024A010797) (RH)

NIMH Grant (R00MH125021) (AW)

NWA-ORC Call (NWA.1160.18.200) (HB, KLH)

ZonMW Top grant (2019/01724/ZONMW) (KLH)

## Author contributions

Conceptualization: AEA, RH, AW, KLH, HB Methodology: AEA, MD

Investigation: AEA, MD Visualization: AEA, MD Supervision: KLH, HDM, HB Writing—original draft: AEA

Writing—review & editing: AEA, KLH, HB, MD, HDM, AW, RH

## Competing interests

H.B. and K.L.-H. are shareholders of Aspect Neuroprofiles BV, which develops physiology-informed prognostic measures for neurodevelopmental disorders. K.L.-H. has filed the patent claim (PCT/NL2019/050167) “Method of determining brain activity”; with priority date 16 March 2018. A.E.-A. is a paid consultant for Aspect Neuroprofiles BV.

