## Supplementary materials for "E/I ratio and net E+I strength are differentially affected across brain disorders"

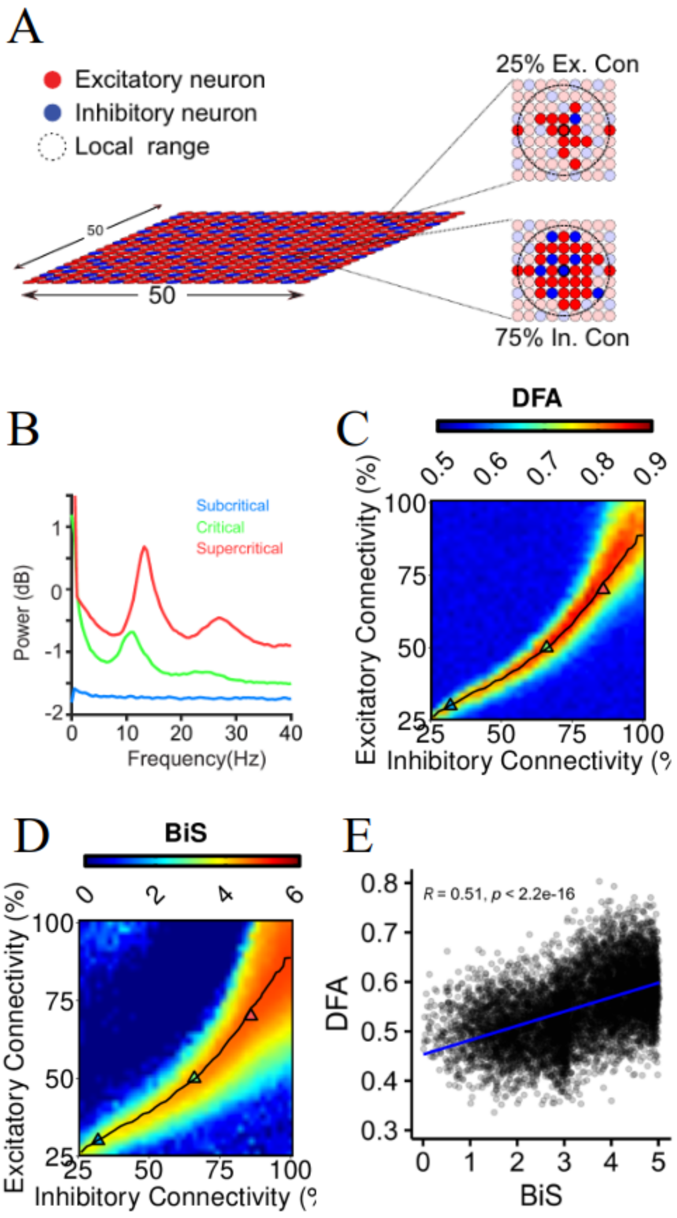


**Figure S1. CROS model: long-range temporal correlations co-occur with bistable oscillations. (A)** CROS model. **(B)** With increasing excitation, we see higher power in alpha oscillations. **(C)** Long-range temporal correlations of alpha oscillations emerge at E/I balance. **(D)** Bistability of oscillations also emerges at E/I balance. **(E)** Criticality and bistability of oscillations are associated.

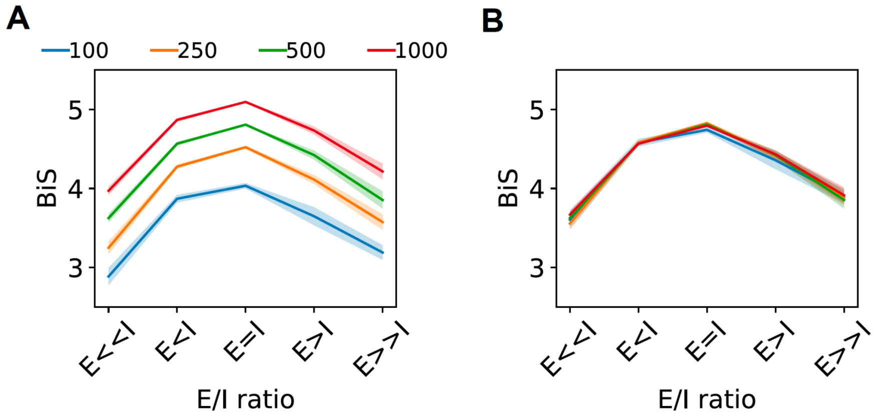


**Figure S2. Bistability depends on the signal length.** We picked 5 networks, with varying levels of E/I ratio: E<<I (exc. conn.=62%, inh. conn.=92%), E<I (exc. conn.=66%, inh. Conn.=88%), E=I (exc. conn.=70%, inh. Conn.=84%), E>I (exc. conn.=72%, inh. Conn.=82%), E>>I (exc. conn.=74%, inh. Conn.=80%), and ran 10 simulations across different network initializations, for a duration of 1000 seconds. We then averaged the bistability index values for each network connectivity combination, and each cropped signal length duration. The averaged bistability values are markedly different when cropping the 1000 seconds signal to shorter lengths (**A**, 50/250/500/1000 seconds) for the original algorithm, but this dependency of BIS value on signal length disappears when using the new formula which corrects for the signal length effects on BIS (**B**). Shaded bands represent ± SEM around the mean.

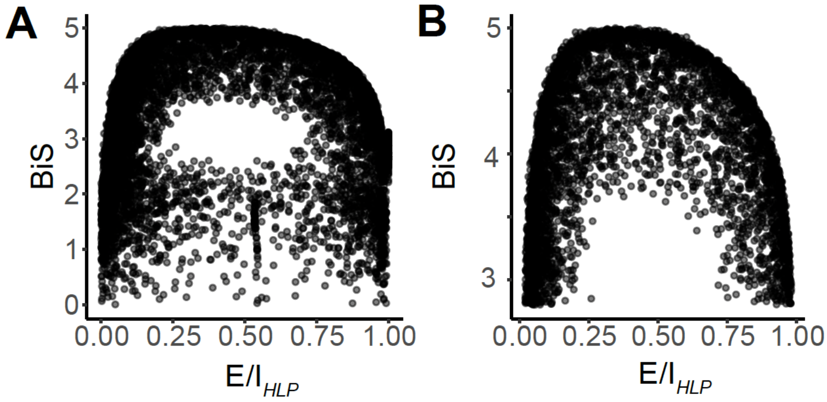


**Figure S3. Inverse-U relationship between bistability and proportion of high-power oscillations helps identify networks where algorithms apply. (A)** $E/I_{HLP}$ ~ BIS relationship for all values. Each dot represents one simulation from the CROS model excitatory/inhibitory connectivity phasespace (Fig. 1A). **(B)** $E/I_{HLP}$ ~ BIS relationship after removing $E/I_{HLP}$ values where BIS<2.5, $E/I_{HLP}$<0.02 or $E/I_{HLP}$>0.98. Each dot represents an individual simulation.

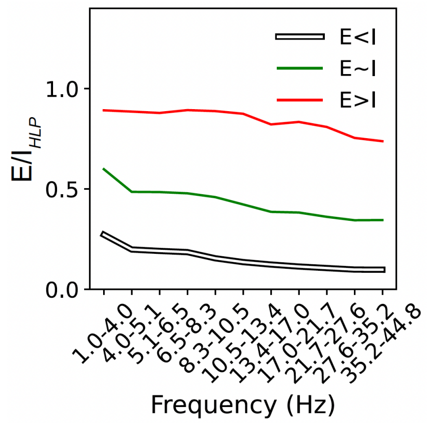


**Figure S4. The proportion of high-power oscillations changes across the spectrum for narrow frequency bins.** In the CROS model, we ran 10 networks for each E/I regime (E<I - *white* triangle in Fig. 1A, E~I - *green* triangle in Fig. 1A, E>I - *red* triangle in Fig. 1A) for 1000 seconds, and computed the E/$I_{HLP}$ values in narrow frequency bands, averaged over 10 runs. Changes in E/I lead to shifts across the spectrum in E/$I_{HLP}$.

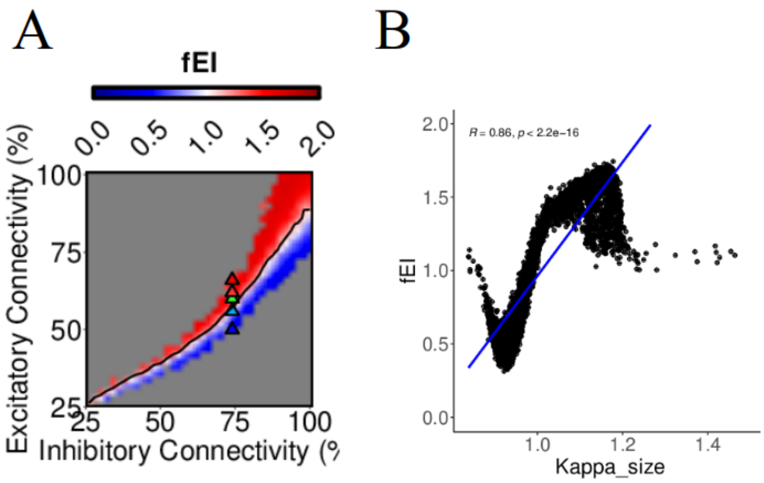


**Figure S5. fE/I can also be used to infer E/I ratio. (A)** fE/I can be used to infer the E/I ratio across the phasespace. fE/I does not apply to networks with DFA<0.6, where the color is gray. **(B)** fE/I can be used to estimate $K_{size}$ of avalanches, which has a value < 1 for inhibition dominated networks, >1 for excitation dominated region, and ~1 at balance.

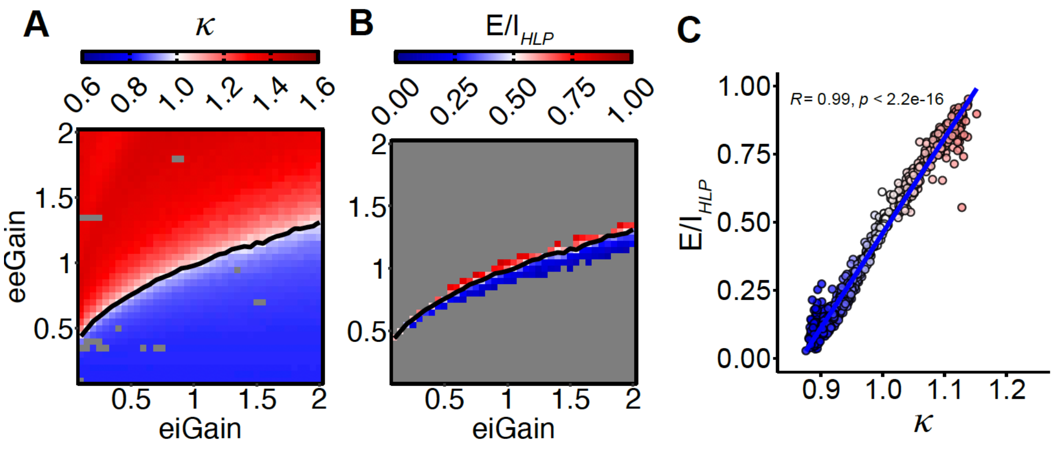


**Figure S6. The proportion of high-power oscillations can also infer E/I ratios in networks modeling synaptic gain changes by neuromodulation. (A)** We modulated the synaptic gain and inhibitory gain of a network close to criticality ($K_{size}$~1), which shifted the dynamics towards a more supercritical regime associated with excitation-dominated networks, ($K_{size}$>1) or a more subcritical regime associated with inhibition-dominated networks. **(B)** $E/I_{HLP}$ could be used to infer the E/I ratio modulation by the changes in synaptic gain, but only for networks with enough evidence of bistability (not in the gray area). **(C)** Where $E/I_{HLP}$ could be estimated, it correlated strongly with $K_{size}$. Each dot represents an individual simulation.

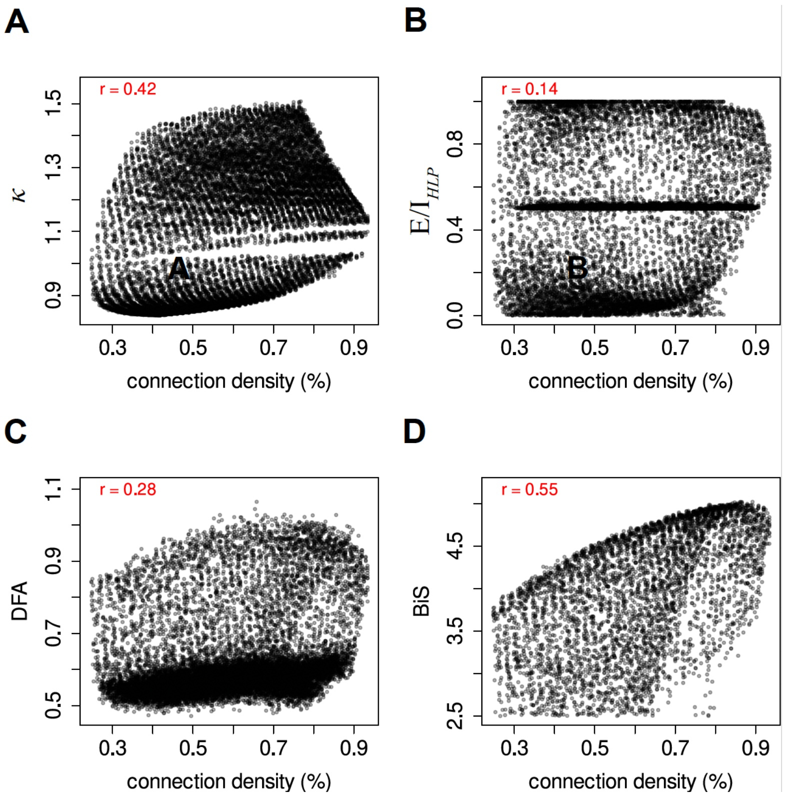


**Figure S7. (A–D) Evaluation of ability of various biomarkers to infer connection density (%).** Each dot represents an individual simulation.

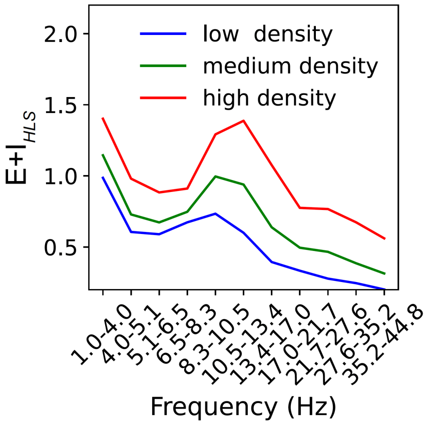


**Figure S8**. **The separation of high- and low-power oscillations changes across the spectrum for narrow frequency bins.** In the CROS model, we ran 10 network initializations for 3 different levels of connectivity density (low connectivity - *blue* triangle in Fig. 2A, medium connectivity - *green* triangle in Fig. 2A, high connectivity - *red* triangle in Fig. 2A) for a duration of 1000 seconds, and computed the $E+I_{HLS}$ values in narrow frequency bands, averaged over 10 runs. Changes in connectivity density lead to shifts across the spectrum in $E+I_{HLS}$.

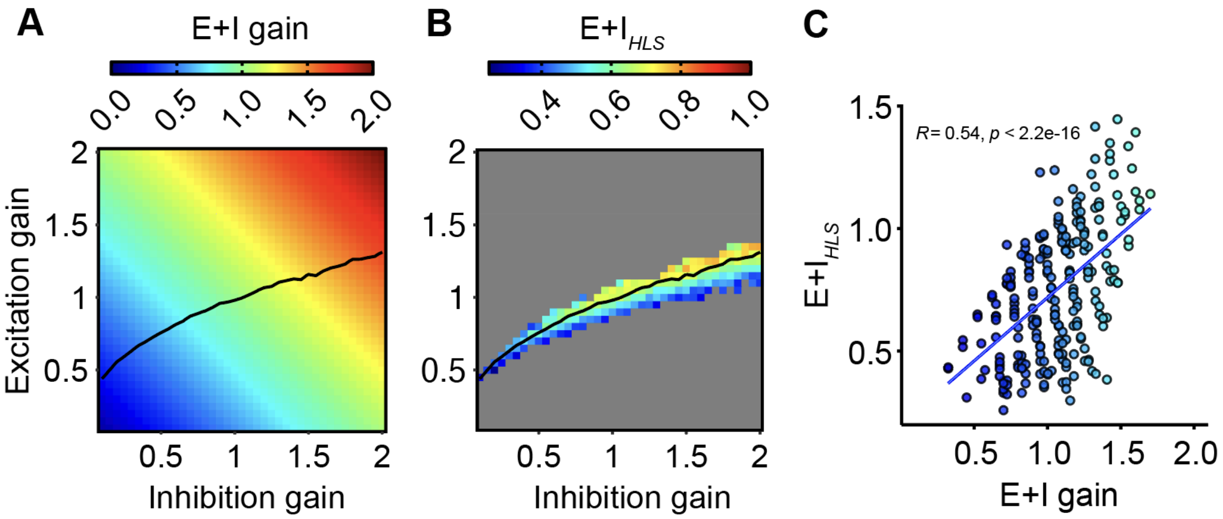


**Figure S9. High-to-low power separation can also be used to infer net E+I strength of synaptic connections in networks modeling neuromodulation of synaptic gain. (A)** We modulated the synaptic gain and inhibitory gain of a network close to criticality ($K_{size}$~1). We estimated the functional E+I density by the sum of the excitatory and inhibitory synaptic gains. **(B,C)** The separation between high- and low-power oscillatory peaks could be used to infer the E+I density. Each dot in C represents an individual simulation.

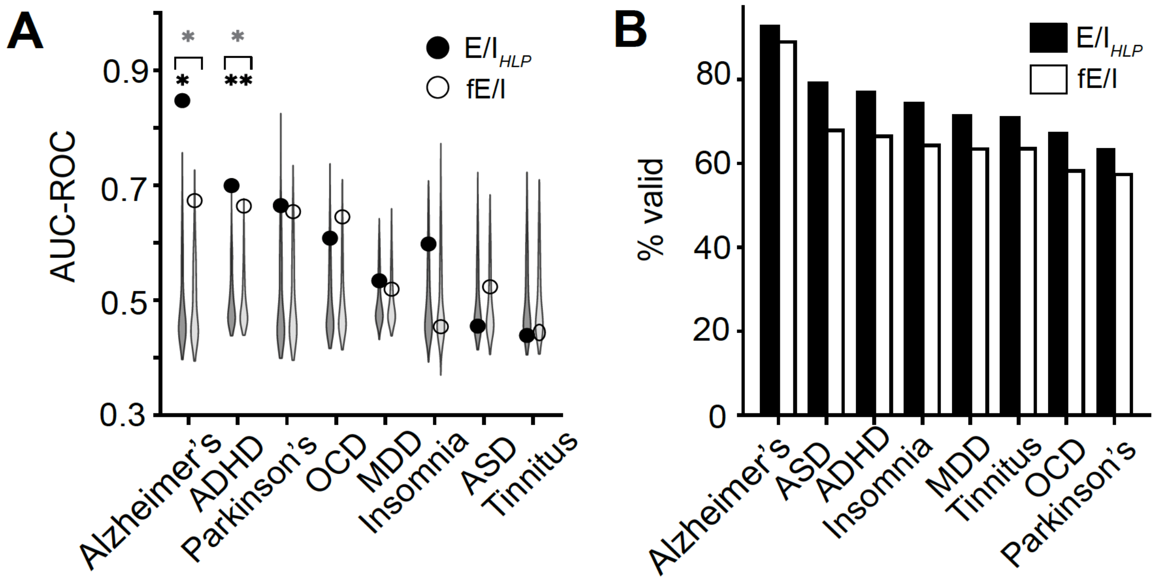


**Figure S10. E/I$_{HLP}$ outperforms fE/I in classification performance for all disorders with significant AUC-ROC and applies to a greater proportion of EEG signals across conditions. (A)** We trained two elastic-net logistic regression models on whole-brain averaged biomarker data for each disorder versus healthy control pair, using either E/I$_{HLP}$ or fE/I features (see *Materials and Methods*). Each model was evaluated with repeated nested cross-validation over 50 folds, and AUC-ROC was computed for each fold. Significance for each disorder was assessed by comparing the average AUC-ROC (black dot for the E/I$_{HLP}$ model and white dot for the fE/I model) across 50 folds against a null distribution generated with 500 random permutations (gray violin plots). Black asterisk reflects significance after FDR ($q$ = 0.05) correction. A paired $t$-test, run across the 50 folds with FDR correction ($q$ = 0.05), was used to assess AUC-ROC differences between the two models, but only for conditions where at least one model showed significantly above-chance AUC-ROC compared to its respective permutation-based null distribution. Gray asterisk reflects significance after FDR ($q$ = 0.05) correction. **(B)** All disorders show a larger percentage of signals where E/I$_{HLP}$ can be applied, compared to fE/I.

| **condition** | ***p*-value** | **significance (FDR corrected)** | **Cohen's *d*** |
|---|---|---|---|
| Alzheimer's | 0.013 | 1 | -0.43 |
| ADHD | $<10^{-5}$ | 1 | 0.74 |
| ASD | $<10^{-20}$ | 1 | 3.25 |

**Table S1. Comparison of AUC-ROC between an E/I$_{HLP}$ model and a E+I$_{HLS}$ model.** We trained two elastic-net logistic regression models on whole-brain averaged biomarker data for each disorder versus healthy control pair, using either E/I$_{HLP}$ or E+I$_{HLS}$ features (see *Materials and Methods*). Each model was evaluated with repeated nested cross-validation over 50 folds, and AUC-ROC was computed for each fold. A paired $t$-test with FDR correction ($q$ = 0.05) was used to assess performance differences between the two models.
